# Calcium-activated chloride channels clamp odor-evoked spike activity in olfactory receptor neurons

**DOI:** 10.1101/282731

**Authors:** Joseph D. Zak, Julien Grimaud, Rong-Chang Li, Chih-Chun Lin, Venkatesh N. Murthy

## Abstract

The calcium-activated chloride channel anoctamin-2 (Ano2) is thought to amplify transduction currents in ORNs, a hypothesis supported by previous studies in dissociated neurons from *Ano2^−/−^* mice. Paradoxically, despite a reduction in transduction currents in *Ano2^−/−^* ORNs, their spike output for odor stimuli may be higher. We examined the role of Ano2 in ORNs in their native environment in freely breathing mice by imaging activity in ORN axons as they arrive in the olfactory bulb glomeruli. Odor-evoked responses in ORN axons of *Ano2^−/−^* mice were consistently larger for a variety of odorants and concentrations. In an open arena, *Ano2^−/−^* mice took longer to approach a localized odor source than wild-type mice, revealing clear olfactory behavioral deficits. Our studies provide the first *in vivo* evidence toward an alternative role for Ano2 in the olfactory transduction cascade, where it may serve as a feedback mechanism to clamp ORN spike output.

## Main Text

Each subtype of olfactory receptor neuron (ORN) converges on a few locations in the olfactory bulb (OB), and therefore serves as a distinct input channel to the brain. ORNs generate electrical signals, in the form of action potentials (spikes), that are interpreted by postsynaptic cells in the OB^1^, including local and projection cells. The series of molecular events that coordinate olfactory transduction and spike generation have been well-delineated, yet much remains unknown about how each individual step in the transduction cascade contributes to overall ORN excitation and output. Specifically, the role of the calcium-activated chloride channel anoctamin-2 (Ano2; also called TMEM16B) remains controversial: many studies point toward its role in massively amplifying ORN transduction currents^2–8^, while paradoxically limiting ORN spike output^9^. In addition, there is conflicting evidence for its importance in olfactory behaviors^8–10^.

Odorants drawn into the nasal cavity bind to odorant receptors (ORs) on the cilia of ORNs. A large number (but not all) of these ORs are G-protein coupled receptors^11–13^, which trigger an intracellular signaling cascade leading to the opening of cyclic nucleotide-gated channels and a net inward flux of Na^+^ and Ca^2+^ ions. The increased intracellular abundance of Ca^2+^ then activates the calcium-activated chloride channel (CaCC) Ano2^9,14,15^. As a result of the elevated intracellular Cl^−^ concentration in ORNs and lower Cl^−^ concentration extracellularly in the nasal epithelium^16–18^, negatively-charged Cl^−^ anions flow outward^16,18^, resulting in an amplification of ORN membrane depolarizations^19^. As much as 90% of the total ORN transduction current may be mediated by Ano2^3,7^, making it a critical component in the sensory transduction pathway that leads to OB input.

Surprisingly, despite its large contribution to the generation of olfactory transduction currents, recent work has suggested that Ano2 is not necessary for odor detection and discrimination^8^. Even the nature of the contribution of Ano2 to ORN activity has become uncertain, since a recent study has suggested that Ano2 may function to limit overall ORN excitability by contributing to a potent depolarization-block of Na^+^ channels^9^. Further investigation of the role that Ano2 plays in olfactory transduction is necessary to understand the precise mechanisms by which odor-evoked excitatory signals are transmitted to the brain.

An additional consideration in the function of Ano2 in olfactory transduction is its expression pattern within ORNs. There is clear evidence that, in addition to being expressed at their cilia^20,11^ within the olfactory epithelium, Ano2 is also abundantly expressed in ORN axons as they terminate in their respective glomeruli^8^. This raises the important possibility that differences between the extracellular Cl^−^ concentration in nasal mucosa/epithelium and the brain may result in Ano2 playing distinct roles in olfactory transduction in different sub-cellular compartments.

How Ano2 contributes to ORN spike output in the native environment of an intact animal, especially at the axon terminals in the OB, have not been addressed. Here, we examine the role of Ano2 in the transmission of odor information to the brain by examining stimulus-evoked responses in ORN terminals in the olfactory bulb of mice with and without Ano2.

## Results

### Glomerular odor maps are unaltered in Ano2-null mice

The axons of ORNs of a common subtype coalesce at a few glomeruli on the surface of the OB. The fasciculation of ORN axon bundles as they enter their respective glomeruli is an activity dependent process^21,22,23^ and could in principle be affected by changes in spontaneous, as well as overall ORN excitability in mice lacking Ano2. Past studies provide conflicting evidence as to whether Ano2 is required for proper targeting of ORNs to glomeruli, with one study reporting that glomerular positioning and number is unaffected in *Ano2*-null mice for two ORN receptor subtypes^8^, while yet another study found an increase in the number of glomeruli incorporating axons of ORNs that express the I7 receptor^9^. We tested whether loss of Ano2 alters functional glomerular maps in the OB.

We crossed heterozygous *Ano2^−/+^* mice (see Methods; Li et al., *In Review*^24^) with a mouse strain that expresses the Ca^2+^ indicator GCaMP3 in all ORNs (OMP-GCaMP)^25^ to obtain two groups of mice, *Ano2^−/−^/OMP-GCaMP3* (KO) and *Ano2^+/+^/OMP-GCaMP3* (WT). First we used wide-field epifluorescence imaging to obtain functional maps of activated glomeruli for seven monomolecular odors by measuring odor-evoked increases in GCaMP fluorescence at ORN axon terminals^26^. These odors (see methods) were selected to activate a diverse number and range of glomeruli on the dorsal surface of the mouse OB^27^.

We compared the total number of glomeruli at each bulb that responded to each odor in our panel for WT and KO mice. We used receiver operating characteristic (ROC) analysis to determine a threshold to define responsive glomeruli, by comparing the response distribution obtained from all glomerulus-odor pairs to a noise distribution obtained from interleaved blank odor trials in each experiment (threshold = 0.005 ∆F/F). There was no significant difference between the number of active glomeruli per OB in WT and KO mice for any of the odors (Figure 1B; p > 0.05, Wilcoxon rank-sum test with Bonferroni multiple comparison correction).

**Figure 1:**
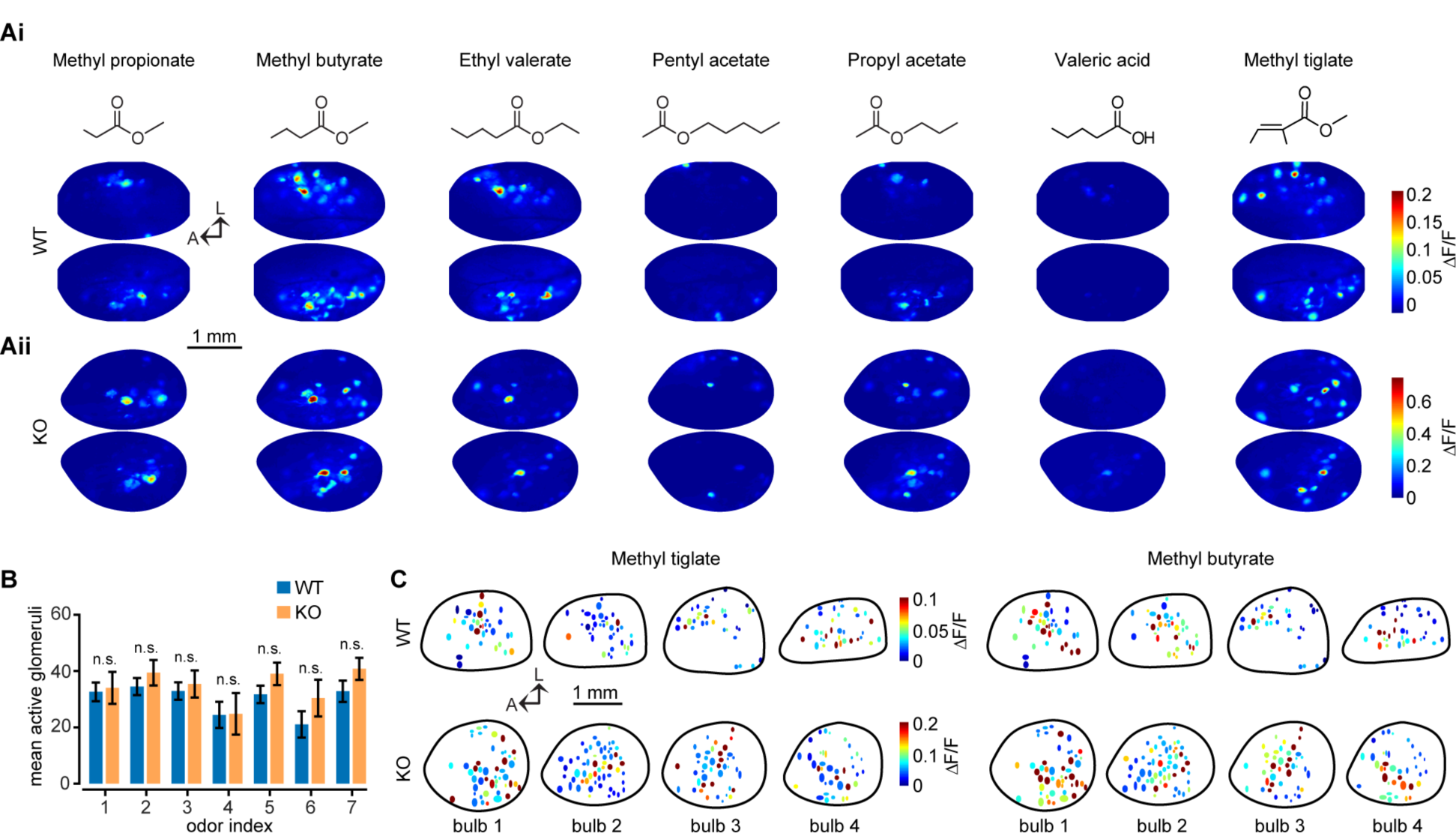
Functional maps of ORN activity. **Ai-Aii**. Example functional maps for seven unique odors in a representative KO (top) and WT (bottom) animal. Molecularstructures of the odors in the panel are depicted above. **B**. Number of glomeruli per bulb (n = 11 WT bulbs, n = 5 KO bulbs) responding to each of the seven odors above ROC threshold (threshold = 0.005 ΔF/F; p> 0.05, Wilcoxon rank-sum test with Bonferroni multiple comparison correction). **C**. Example functional maps from four WT and KO bulbs each for two different, but related odors.

Although our experiments were not designed to identify specific glomeruli, we can nevertheless discount large scale glomerular duplication in *Ano2* KO animals, for example due to a loss of targeting specificity and mixing of ORN axons within individual glomeruli^28^. We also cannot discount small-scale redistribution of glomerular positioning, which is unlikely to exceed expected animal-to-animal variance described previously in control mice^26,28^. Together, these results indicate ORN maps are largely conserved in KO mice and confirm previous anatomical studies^8^.

### ORN input to glomeruli is enhanced in Ano2-null mice

We next asked whether odor-evoked responses in individual glomeruli were altered in either magnitude or duration in KO mice. From six WT mice (n = 9 bulbs) and three KO mice (n = 5 bulbs), we identified 34.44 ± 3.48 and 43.60 ± 3.93 glomeruli per bulb, respectively (299 total glomeruli in WT mice and 218 total glomeruli in KO mice). The 50 largest responses for each odor are shown in Figure 2A. Our analysis revealed significantly larger odor-evoked responses in ORNs of KO mice than WT mice (n = 2093 WT and 1526 KO glomerulus-odor pairs, p <0.001 Kolmogorov–Smirnov test; Figure 2B). This difference was observed both when responses of each glomerulus was averaged across all seven odors (n = 299 WT glomeruli and 218 KO glomeruli, p < 0.001, Kolmogorov–Smirnov test), and when responses of all glomeruli for each odor were averaged (Figure 2C-E).

**Figure 2:**
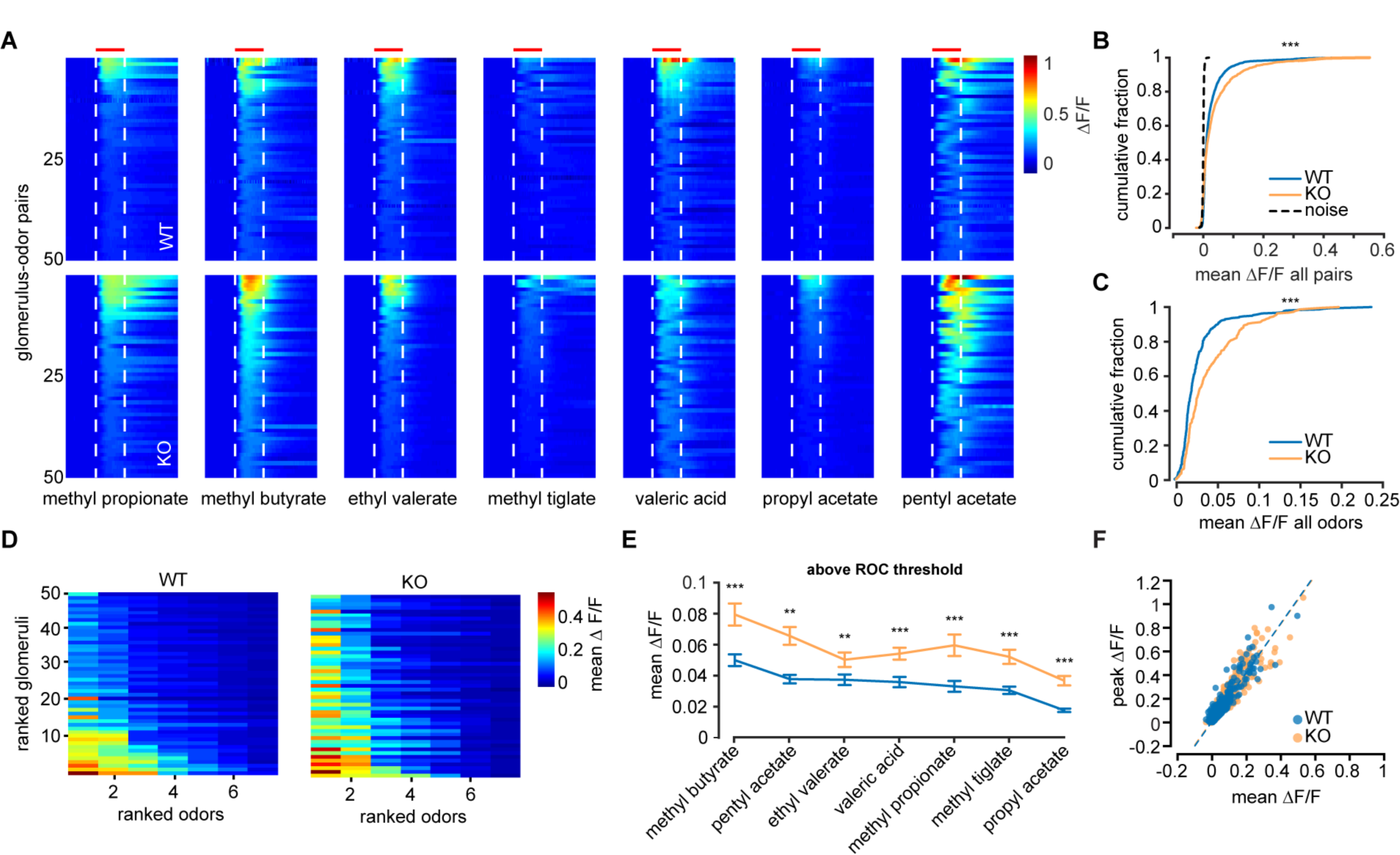
Odor responses in *Ano2*-null mice. **A**. The 50 largest odor-evoked Ca^2+^ signals across all animals for each of seven odors in bothgroups. Dashed lines and red bar indicate odor delivery period. Data are sorted by the largest mean response during odor delivery. **B**. Cumulative distribution of the meanCa^2+^ response in the odor period across all glomerulus-odor pairs (n = 2093 WT and 1526 KO glomerulus-odor pairs, p < 0.001 Kolmogorov–Smirnov test).**C**. Cumulative distribution of the mean Ca^2+^ response across all odors at each glomerulus (n = 299 WT glomeruli and 218 KO glomeruli,p < 0.001, Kolmogorov–Smirnov test). **D**. The top 50 glomeruli rankedby mean response across all odors and further ranked by individual odor responses for WT(right) and KO (left) animals. **E**. Mean response of all glomeruli responding aboveROC threshold (threshold = 0.005 ΔF/F, Wilcoxon rank-sum test with Bonferroni correction, *p < 0.05, **p<0.01, ***p<0.001). **F**. Scatter plot of the peak response as a function of the mean response for all glomerulus-odor pairs. Linear regression for each group plotted as dashed line (slope = 2.079, R^2^ = 0.887 for WT and slope = 2.023, R^2^ = 0.881 for KO)

We next asked whether Ca^2+^ response kinetics in ORNs were different in WT and KO mice. First, we compared the peak response with the average response over the entire period (5 s) of odor presentation. Our analysis revealed a strong linear correlation between the two parameters in both WT and KO mice, as well as nearly identical slopes of regression (slope = 2.079, R^2^ = 0.887 for WT and slope = 2.023, R^2^ = 0.881 for KO, Figure 2F). This result indicates that the larger odor responses in KO animals were due to scaling up, rather than a temporal redistribution, of spikes in ORNs. We also used principal component analysis (PCA) to compare the response time course of each group (Supplemental Figure 1) and found no significant differences in the Ca^2+^ responses of WT and KO animals.

Our data provide evidence that loss of Ano2 results in enhanced ORN input to OB glomeruli without impacting the overall response time course or duration. We also observed that the number of glomeruli responding to an odor was similar in WT and KO animals. These results are consistent with a scaling mechanism, where Ano2 may function as a negative feedback on ORN excitability and limit the number of action potentials generated in response to odor stimulation.

### Loss of Ano2 does not impact respiration

It is possible that the larger responses to odorants in *Ano2* KO mice is due to faster respiration rates and temporal summation of responses^29^. Other ANO isoforms are expressed in smooth muscle tissue and may regulate the excitability of the diaphragm and the airway^30,31,32,33^, thereby altering normal breathing rhythms in KO mice. We recorded the respiration rate of WT, KO, and C57BL/6J mice using a thermocouple placed in front of the animal’s nose, under anesthesia conditions consistent with our previous experiments. We first validated the reliability of the external thermocouple for respiration tracking by comparing it to a well-established method, intranasal cannulas (Supplemental Figure 2). Upon validation, we chose to record respiration using the non-invasive thermocouple to mitigate any effects of damaging the nasal cavity through cannula implantation.

Robust respiration signals could be recorded in anesthetized mice (n = 6 WT, 7 KO, 3 C57BL/6J; Figure 3A for example traces). We included C57BL/6J animals in our comparison to rule out any genetic background effects^34^. Despite slight animal-to-animal variability, we found that across all groups, there was no statistical difference in the overall respiration frequency (mean frequency: 1.60 ± 0.17Hz WT, 1.86 ± 0.23Hz KO, 1.77 ± 0.19Hz C57BL/6J, p = 0.71 Kruskal-Wallis test; Figure 3C). The small, statistically insignificant increase in respiration rate we observed in KO mice (16.3% increase) is insufficient to account for the larger Ca^2+^ responses obtained in our previous imaging experiments (p < 0.001, Kolmogorov–Smirnov test).

**Figure 3:**
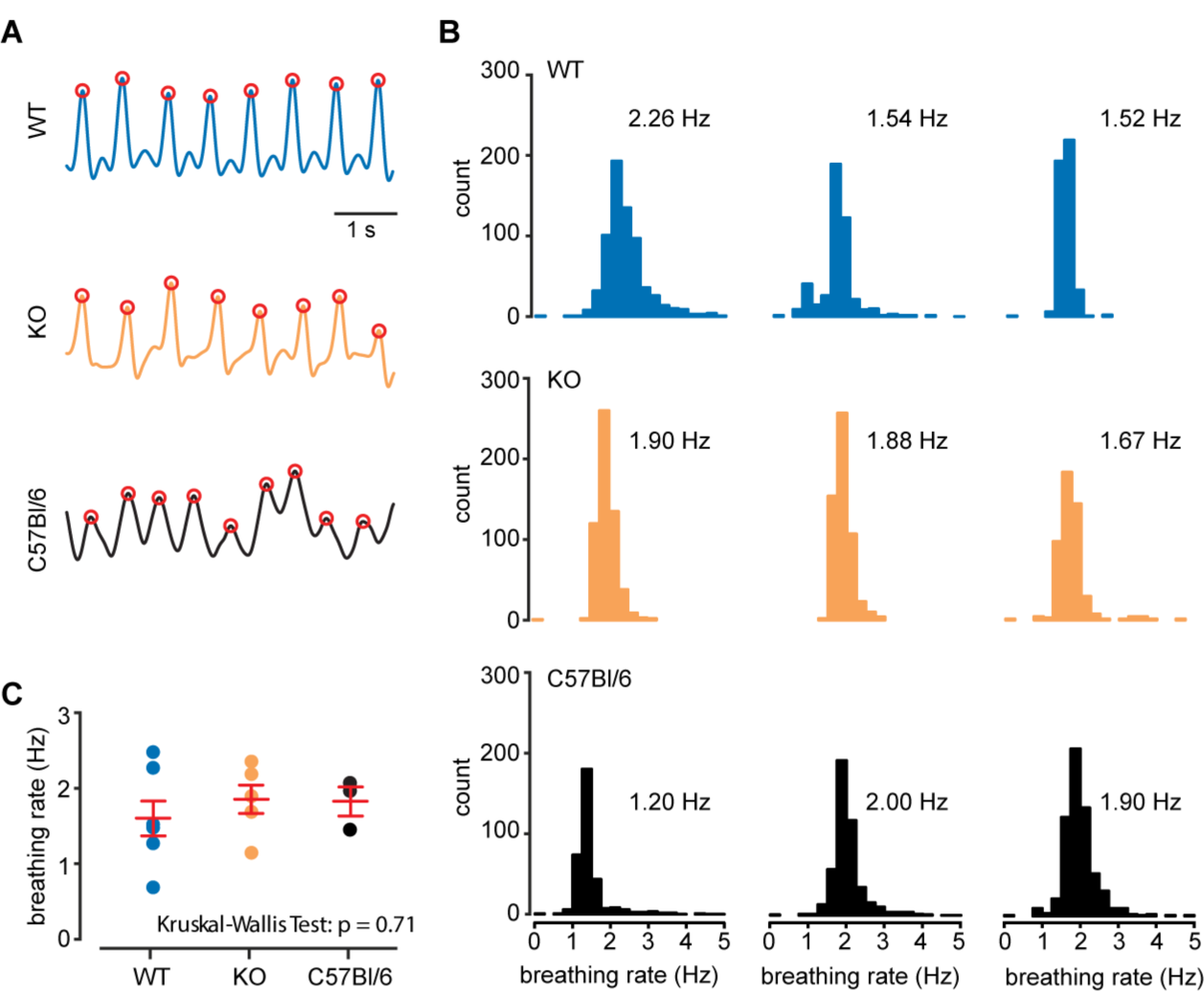
Loss of Ano2 does not impact respiration rate. **A**. Example respiration traces from a WT, KO and C57BL/6J animal recorded with a thermocouple placed near the animal’s nose (see Supplementary Figure 2 for technique validation). **B**. Histograms of the instantaneous respiration frequency in a 5-minute window from three representative animals from each group. Mean instantaneous frequency is displayed next to each plot. **C**. Mean instantaneous frequency for all animals in each group. Red bars denote mean and standard error across all animals (p = 0.71, Kruskal-Wallis test).

We conclude that anesthetized KO animals do not breathe with increased frequency and that the larger Ca^2+^ signals we observed are not due to enhanced respiration rate.

### Multiphoton imaging in Ano2-null mice

Due to the low resting fluorescence of GCaMP3 we were unable to identify glomeruli that did not respond to at least one of the seven odors in our panel using an epifluorescence microscope. We used multiphoton microscopy to overcome this limitation and were able to visualize all glomeruli, independent of their responsiveness (Figure 4A). We also expanded our odor panel size to 15 odors to activate a wider range of glomeruli. The optical sectioning facilitated by multiphoton microscopy also allowed us to exclude any effects arising from the activity of *en passant* axons that could be detected using our epifluorescence setup. For example, strongly excited ORNs passing over an otherwise inactive glomerulus will give the appearance of Ca^2+^ activity at the glomerulus below.

**Figure 4:**
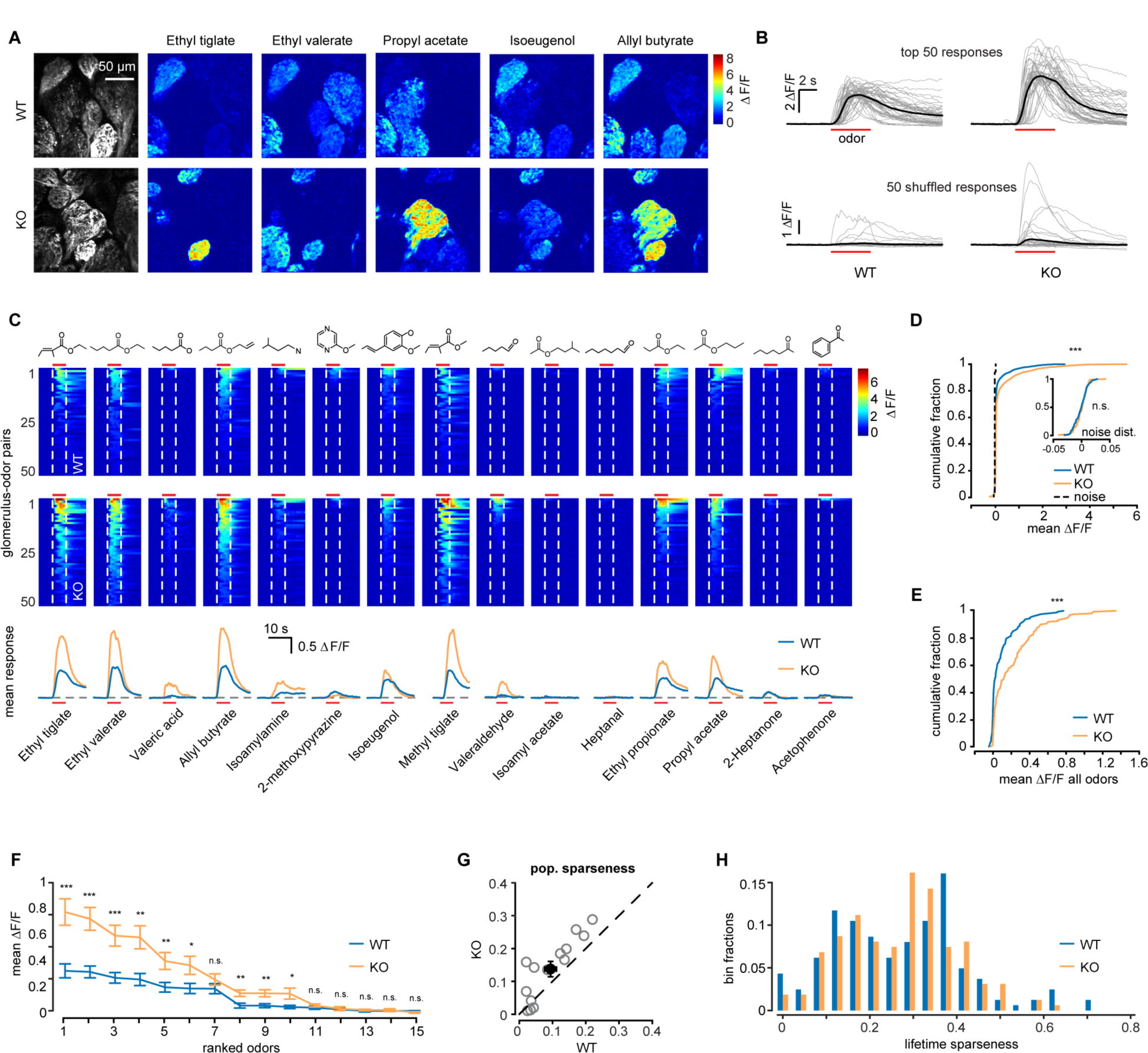
Calcium responses in ORNs measured with multiphoton microscopy. **A**.Example multiphoton-acquired images of glomeruli from an example WT and KO animal, as well as example ΔF/F responses for five selected odors. Mean images from 20 frames preceding and during the odor delivery period were used. **B**. Traces of the 50 largest (top) Ca^2+^ responses for WT and KO animals across all odors and 50randomly selected responses (bottom). **C**. The 50 largest odor-evoked Ca^2+^ signals across all animals for each of 15 odors. Dashed line denotes odor onset and offset and red bar indicates odor duration. Molecular structures are depicted above. Bottom, mean response time course for each odor. **D**. Cumulative distribution ofthe mean Ca^2+^ response in the odor period across all glomerulus-odor pairs (n = 2430 WT and 2415 KO glomerulus-odor pairs, p < 0.001, Kolmogorov–Smirnov test). Inset, distribution of blank odor trial responses used to determine ROC threshold (threshold = 0.02 n = 162 WT and 161 KO glomeruli, p > 0.05, Kolmogorov–Smirnov test). **E**. Cumulative distribution of the mean Ca^2+^ response across all odors at each glomerulus (n = 162 WTand 161 KO glomeruli, p < 0.001, Kolmogorov–Smirnov test). **F.** Mean response of all glomeruli responding above threshold for each odor (Wilcoxon rank-sum test with Bonferroni correction, *p < 0.05, **p<0.01, ***p<0.001).**G**. Scatter plot of population sparseness for each odor. Mean across all odors is the filled black circle (mean sparseness =0.09 ± 0.02 in WT and 0.13 ± 0.02 in KO, p = 0.002, Wilcoxon sign-rank test). **H**. Histogram of lifetime sparseness across all glomeruli (mean sparseness = 0.28 ±0.01 in WT and 0.28 ± 0.01 in KO, p = 0.89, Wilcoxon rank-sum test.

Across 162 WT and 161 KO glomeruli from five animals each, we found that ORNs in KO animals responded with significantly larger Ca^2+^ transients, whether comparing individual glomerulus-odor pairs (n = 2430 WT and 2415 KO glomerulus-odor pairs, p < 0.001, Kolmogorov–Smirnov test; Figure 4D), or mean response across all odors (n = 162 WT and 161 KO glomeruli, p < 0.001, Kolmogorov–Smirnov test; Figure 4E). The 50 largest overall responses for each group are displayed in Figure 4B (mean of all responses above threshold:38.8 ± 0.02% ∆F/F in WT and 57.5 ± 0.03% ∆F/F in KO). When considering each odor individually, in 9 out of 15 odors we observed significantly larger Ca^2+^ responses in KO animals (Figure 4H, Wilcoxon rank-sum test with Bonferroni correction). We again examined the time course of the Ca^2+^ responses in each group using PCA (Supplemental Figure 3) and found no difference between WT and KO animals.

We next compared sparsity of glomerular responses. First, we calculated the population sparseness (see Methods) to compare the fraction of activated glomeruli across all animals for each odor. We found that a larger fraction of glomeruli responded in KO animals (population sparseness measure: 0.09 ± 0.02 in WT and 0.13 ± 0.02 in KO, p = 0.002, Wilcoxon sign-rank test, Figure 4G). When we compared lifetime sparseness, which quantifies the extent to which a given glomerulus responds to different odor stimuli, we found no difference (lifetime sparseness measure: 0.28 ± 0.01 in WT and 0.28 ± 0.01 in KO, p = 0.89, Wilcoxon rank-sum test, Figure 4I). Together these results indicate that ORNs in KO animals are indeed more sensitive to odor stimulation, but the breadth of their odor tuning is unchanged. Furthermore, the fact that we observed no difference in odor tuning further argues against the possibility that glomeruli in KO animals receive heterogeneous innervation from multiple ORN subtypes. We also found no evidence for heterogeneous responses within individual glomerular regions of interest.

### ORNs are more strongly excited by odors across a range of concentrations

What accounts for the larger fraction of activated glomeruli in KO animals? One possibility is that the signal-to-noise ratio afforded by multiphoton microscopy allowed us to identify weak responses arising from odor-receptor binding in KO animals that are sub-threshold for Ca^2+^ signal generation in WT animals. Conversely, given our previous results, another explanation for the increased Ca^2+^ signal magnitude in KO mice is that in response to high odor concentrations, ORNs are able to maintain firing due to a reduction in depolarization induced Na^+^ inactivation driven by the amplifying current through Ano2. However, at low odor concentrations, ORNs in KO animals may have weaker responses than ORNs in WT animals, since it has been shown that current amplification through Ano2 is most potent close to detection threshold (Li et al., unpublished). We next investigated whether ORNs in KO animals are more responsive to odors at different concentrations.

We used air dilution to alter odor concentration over four orders of magnitude for two odors, Ethyl valerate and Allyl butyrate, and decreased the odor delivery time to two seconds to prevent saturation of ORN responses at the highest concentrations. The relative concentration of each odor experienced by the mouse was verified using a photoionization detector and odor concentrations were normalized to the lowest dilution (see Methods; Supplemental Figure 4). At the strongest odor concentrations, glomerular ORN responses were again enhanced in KO animals for both odors, as well as a mixture of the two (Figure 5B-D, Wilcoxon rank-sum test). However, somewhat surprisingly, at low concentrations our analysis revealed no pair-wise differences. We were unable to compare the response amplitude of ORNs at their individual detection thresholds due to an inability to identify specific subtypes of ORNs (expressing a particular odorant receptor) across animals. We also note that Ca^2+^ signals in ORN axons report spike output rather than transduction current amplitudes. Our results thereby indicate that while Ano2 may play a role in shunting ORN excitation following strong odor stimulation, it does not limit ORN activity following weak odor input.

**Figure 5:**
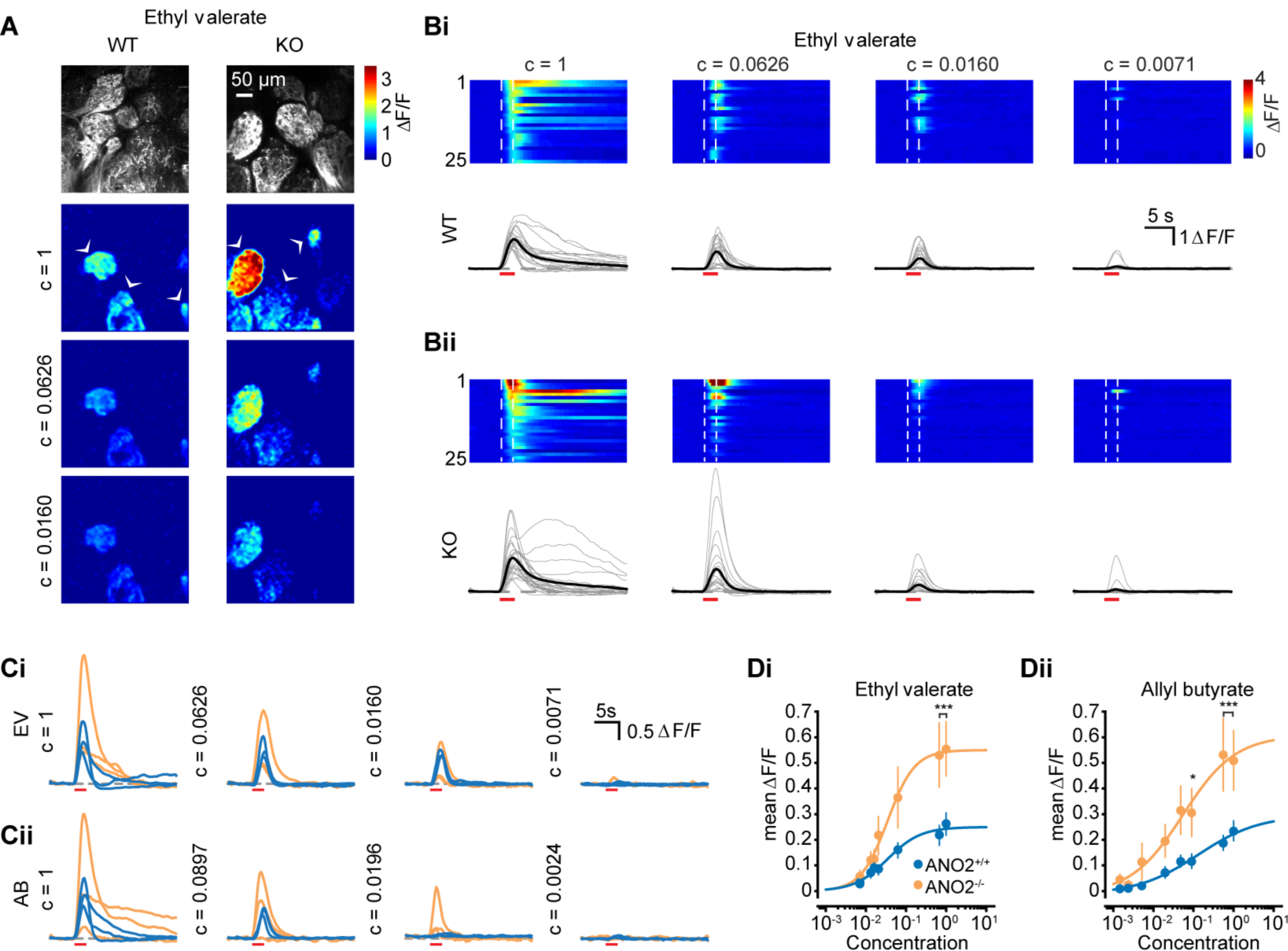
ORN responses in Ano2-null mice are enhanced at high odor concentrations.**A**. Example multiphoton-acquired images of glomeruli and ΔF/F responses at three odor concentrations. Odor concentrations were normalized to the highest concentration (~10% v/v) using a photoionization detector (see Supplemental Figure 4).**B**. Examples of the 25 largest responses to the highest concentration of Ethyl valerate followed through four other concentrations. Individual traces are displayed below.Dashed line denotes odor onset and offset, red bar indicates odor period. **C**. Example traces from three glomeruli in part A, identified by arrowheads followed through four concentrations of Ethyl valerate (EV; top) and Allyl butyrate (AB; bottom) **Di-ii**. Mean response (filled circles) at eight odor concentrations for both odors and sigmoidal fit to each (Wilcoxon rank-sum test with Bonferroni correction, *p <0.05,**p<0.01, ***p<0.001).

### Ano2 deletion increases latency to odor localization

Does increased ORN excitability alter odor detection capabilities of KO animals? Recent studies provide evidence that *Ano2*-null mice exhibit a greater latency to uncover a hidden food-object^9,10^, while another study was unable to find any difference in odor detection and discrimination using a learned behavior^8^. We decided to study innate odor-driven investigation to assess whether KO animals displayed any sensory deficit independent of learned behavior.

We investigated the latency to explore odors as an indicator of how easily mice can detect odors^9,10^. To minimize experimenter-induced biases, our automated experimental apparatus consisted of a 56 cm diameter circular arena with four air inlets equally spaced around its circumference, as well as a vacuum in its center to balance air inflow and outflow. Under infrared illumination, mice were allowed to explore the arena space for 10 minutes, after which odorized air was delivered through one of the inlets. We then measured the latency of each animal to investigate the odor source as determined by the animal approaching the odorized air inlet within 1cm.

We used the known appetitive odor peanut oil^35^ diluted to the same concentration as we used in our imaging experiments (1% in mineral oil). Consistent with previous reports^9,10^, across 8 WT and 11 KO animals, we found that KO animals required significantly more time to locate the odor source (Figure 6A, p = 0.003, Wilcoxon rank-sum test). Because the odor onset occurred independently of the animal location in the arena, we calculated the initial starting distance from the odor source and observed no differences in their mean positions (Figure 6B, p > 0.05, Wilcoxon rank-sum test). At the same time, we observed no differences in the locomotor activity of KO animals as measured by their mean velocity both prior to and following odor delivery (Figure 6C-D, p > 0.05, Wilcoxon rank-sum test). These behavioral data suggest a puzzling dissociation between the increased responses to odorants in KO animals and the longer latency to locate the source of an appetitive odor.

**Figure 6:**
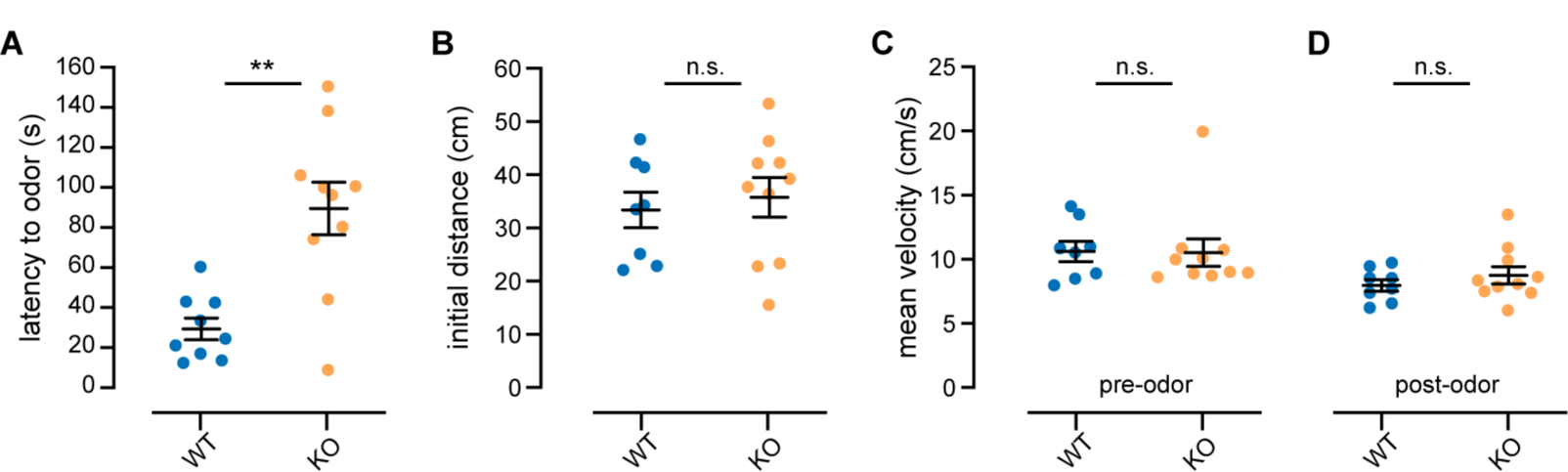
Odor localization latency is increased in Ano2-null mice. **A**. Time latency for mice to locate the source of odorized air carrying a 1% dilution ofpeanut oil (n = 8 WT and 11 KO animals, p = 0.003, Wilcoxon rank-sum test). **B**. Plot of the initial starting distance across all animals used (p > 0.05,Wilcoxon rank-sum test). **C-D**. Plot of the mean velocity of each animal prior toand following odor onset (p > 0.05, Wilcoxon rank-sum test).

## Discussion

Our study presents direct evidence in freely-breathing mice that Ano2, despite its role in amplifying transduction currents in ORNs, limits their overall excitation and input to the OB *in vivo*. Our results are in agreement with recent *in vitro* measurements of ORN spike output in *Ano2*-null mice and further point towards a dual functionality of Ano2 in ORN excitability whereby it both amplifies transduction currents and limits spike output.

### Glomerular maps and respiration

Loss of Ano2 in mice could lead to more general changes that might be confounding factors that undermine conclusions about sensory transduction and coding. First, absence of Ano2 might alter the anatomical organization of glomerular maps in the OB. In particular, spontaneous activity in ORNs is known to play an important role in ORN fasciculation and glomerular emergence^21–23^. Any differences in spontaneous activity between WT and *Ano2* KO mice might lead to disorganized glomerular organization and odor representation. Our results argue against a broad topographical reorganization of ORN inputs to the OB in KO animals based on two factors. First, we found that the number of dorsal glomeruli responding above threshold to a given odor was unchanged and second, we found no difference in the lifetime sparseness of individual glomeruli from WT and KO animals. Furthermore, glomeruli in KO animals do not appear to receive heterogeneous ORN innervation since responding glomeruli were invariably homogeneous.

A second factor that might affect the data on glomerular imaging is the respiration rate. Faster respiration may lead to larger Ca^2+^ signals because of slow time course of axonal Ca^2+^ as well as indicator kinetics. Direct measurement of respiration, however, dispelled this concern – we found no significant change in respiration rate in *Ano2* KO animals. On a methodological note, we also demonstrated that an externally-placed, non-invasive thermocouple is a reliable method to record and measure breathing responses in anesthetized mice. While this method could be valuable for experiments in anesthetized animals, we note that it rapidly loses fidelity when breathing rate increases, as in awake animals (Supplementary Figure 2E,F).

### Larger odor-evoked responses in KO mice

The major finding of our study is that the magnitude of the ORN Ca^2+^ responses following odor stimulation was larger in KO animals, with no observable change in the overall response duration. This was confirmed in two different modes of imaging – widefield microscopy that allowed larger regions to be imaged at lower resolution, and multiphoton microscopy that offered excellent optical sectioning and signal-to-noise ratio. We systematically varied the concentration of two different odorants and found that responses in ORNs from KO animals were consistently larger at most concentrations. Interestingly, at lower concentrations of the two odors, the response amplitudes were similar between both groups. This result suggests that for low odor concentrations, ORN transduction currents may remain sufficiently modest, and limits further amplification through Ano2.

Biophysical studies *in vitro*, however, indicate that Ano2 currents are activated even for weak stimuli^10,36^. *In vitro* preparations allow for titration of odor concentrations for each neuron, thus allowing careful analysis of transduction currents in different response regimes, including threshold and sub-threshold responses. Perhaps glomerular imaging does not have enough sensitivity to detect responses to low concentrations of odors, and potential differences between WT and KO mice were missed. *In vivo*, threshold odor concentrations are likely to activate only a subset ORNs due to their dispersion throughout the nasal epithelium^37^. The population imaging approach used in our studies further decreases the likelihood of detecting low-level signals because any responses arising a small number of ORNs are averaged across all ORNs that terminate at a given glomerulus. An alternative approach may include sparsely labeling ORNs with Ca^2+^ indicators to allow for recordings of individual optically isolated axon terminals; however, to date there is no reliable method available for such an approach.

Another key result is that responses saturated at lower amplitudes in WT than in KO glomeruli, suggesting that the presence of Ano2 had a “clamping” effect, and its absence loosens the clamp to allow greater activity. Our findings and work of others^9^, suggest that in a high odor concentration regime, Ano2 may function as a feedback mechanism to limit the number of spikes generated by ORNs following odor stimulation. The proposed mechanism of action operates through a potent depolarization-induced inactivation of Na^+^ channels following transduction current amplification by Ano2^9^. A potential caveat is that the larger responses we observe in KO mice are the result of lower resting fluorescence due to a decrease in spontaneous ORN activity at some glomeruli^9^, thereby increasing the dynamic range available for Ca^2+^ indicator activity. Our experiments used odor concentrations that are generally thought to be sub-saturating for odor evoked responses in ORNs; however, it remains possible that at these concentrations, the largest signals observed in WT animals exceeded the range of our indicator due to a greater basal Ca^2+^ tone. Our results here argue against this possibility since the only observable differences occurred in response to strong odor stimulation – larger responses in ANO KO mice would not be observed if ORN responses in WT were saturated at these higher concentrations.

It is also possible that Ano2 alters neural excitability in other ways, especially since Ano2 is expressed in ORN terminals^8,38^. For example, in the thalamocortical^39^ system Ano2 functions to suppress neural excitability by enhancing the magnitude of action potential after-hyperpolarization. In the hippocampus^40^, Ano2 decreases the duration of individual action potentials by relying on a chloride gradient that favors outward membrane currents close to resting potential^41^. Although the chloride gradient in the nasal epithelium favors inward currents at near resting potentials^16,18^, the presumed chloride gradient at ORN axonal terminals could yield outward currents through Ano2, triggered by inward flux of Ca^2+^ during action potentials. A reduction in ORN transduction currents at the olfactory cilia of *Ano2*-null animals could be offset by a reduction in the action potential after-hyperpolarization in the axonal compartments of ORNs.

Independent of the mechanisms involved, Ano2 seems to functionally compress the dynamic range of odor responses in individual ORNs by leaving weak responses unaffected (or enhancing them) and truncating the magnitude of responses to strong odors.

### Role of Ano2 in olfactory coding

Our data suggests that currents through Ano2 may serve as a negative feedback mechanism to prevent excessive activation at higher concentrations. It remains possible that at low to moderate concentrations, Ano2 may act to amplify sensory signals and affect activity in ways undetected by our measurements. For instance, the latency to first spike (from the onset of inhalation) could be shorter in WT ORNs because of the amplification by Ano2. Such changes in timing could play an important role in odor coding^42,43,44^, but our imaging methods may not have the temporal resolution or sensitivity to detect such differences in latency. It is apparent that simply scaling up the activity in ORNs is insufficient enhance odor detection and may instead have deleterious effects.

Another potential role of Ano2 may arise from differences in expression of Ano2 in ORNs of a common subtype. Through varying expression levels, ORNs projecting to the same glomerulus could de-correlate their firing patterns in response to the same stimulus by shunting their spike output at different levels and thereby increasing their information carrying capacity as a population. Past studies demonstrate that intrinsic biophysical diversity between sister mitral cells functionally reduces correlations in their spike output^45^ and these observations are consistent in other systems including ganglion cells^46^ and M1 type ganglion cell photoreceptors^47^ in the retina. In our mouse line, all ORNs were labeled with GCaMP3 and we were therefore unable to study heterogeneity at the single cell level. However, future studies may seek to record from a small number of ORNs projecting to the same glomerulus to determine whether Ano2 plays a role in de-correlating their spike output. Furthermore, a decreased information carrying capacity of ORNs in *Ano2*-null mice provides a potential explanation for the odor localization deficits observed in this study and others^9,10^. We observed that *ANO2* KO animals, despite having larger odor-evoked ORN responses, require longer to locate a relatively low-concentration odor source. This result suggests that deletion of Ano2 does not simply scale up the sensitivity of ORNs, but rather, results in a fundamental reformatting of how odor information is transmitted to the brain. At present it is not clear how odor information is restructured in ORNs of KO animals, or if dysfunction in odor information processing is further compounded by downstream neurons.

## Materials and Methods

### Animal Care, General Statements

C57BL/6J, *Ano2^+/+^*, *Ano2^−/−^,* and OMP-GCaMP3 mice were used in this study. The age of all animals at the time of the experiments was two to six months. All mice used in this study were housed in an inverted 12-hour light cycle and fed *ad libitum*. All the experiments were performed in accordance with the guidelines set by the National Institutes of Health and approved by the Institutional Animal Care and Use Committee at Harvard University.

### Animal Generation

The *Anoctomin-2* knock-out (*Ano2^−/−^*) mouse line was obtained from the PBmice project of Fudan University (http://idm.fudan.edu.cn/PBmice). The line was generated with a piggyBac transposon system^34^. A donor plasmid carrying the terminal sequences required for transposition flanking a fluorescence protein gene driven by actin promoter (Act-red fluorescent protein (RFP), to facilitate selection), and a helper plasmid carrying the PB transposase (PBase) were injected into the pronucleus of mouse embryonic stem cells. The terminal sequences and Act-RFP were integrated to the genome and were stably transmitted through germline, while the plasmid carrying PBase was lost in subsequent progenies. The mouse line with piggyBac transposon inserted into intron 14 of *Ano2* was then identified.

### In vivo imaging

#### Surgery

Adult mice were anesthetized with an intraperitoneal injection ketamine and xylazine (100 and 10 mg/kg, respectively) and eyes were covered with petroleum jelly. The scalp was shaved and opened. After thorough cleaning and drying, the exposed skull was gently scratched with a blade, and a titanium custom-made headplate was glued on the scratches. The cranial bones over the OBs were then removed using a 3 mm diameter biopsy punch (Integra Miltex). The surface of the brain was cleared of debris and a glass coverslip was glued into the vacated cavity in the skull. Dental cement (Jet Repair, Lang Dental) was used to cover the headplate and form a well around the cranial window. Mice were allowed to recover for at least three days. Prior to each imaging session, animals were anesthetized with a mixture of ketamine and xylazine (90% of dose used for surgery) and body temperature was maintained at 37°C by a heating pad.

#### Epifluoresence

Two photo lenses coupled front to front were used to image the OB surface onto the sensor of a CMOS camera (DFK 23GPO31, The Imaging Source GmbH). Images (960 × 600 pixels) were acquired at 8-bit resolution and 8 frames/s. Data from the camera were recorded to the computer via data acquisition hardware (National instruments) and custom software in Labview. A blue LED (CBT-90, Luminus) with a maximum output of 1.65mW/mm^2^ was used for excitation.

#### Multiphoton

A custom-built two-photon microscope was used for in vivo imaging. Fluorophores were excited and imaged with a water immersion objective (20×, 0.95 NA, Olympus) at 920nm using a Ti:Sapphire laser (Mai Tai HP, Spectra-Physics). Images were acquired at 16-bit resolution and 4 frames/s. The pixel size was 1.218µm, and fields of view were typically 365 x 365µm. The point-spread function of the microscope was measured to be 0.51 x 0.48 x 2.12 µm. Image acquisition and scanning were controlled by custom-written software in Labview. Emitted light was routed through two dichroic mirrors (680dcxr, Chroma and FF555-Di02, Semrock) and collected by a photomultiplier tube (R3896, Hamamatsu) using filters in the 500-550 nm range (FF01-525/50, Semrock).

#### Odor stimulation

Monomolecular odorants (Sigma) were used as stimuli and delivered by custom-built 8 channel (epifluorescence experiments) or 16 channel (2-photon experiments) olfactometer controlled by custom-written software in Labview (National Instruments)^27^. Odorants were maintained at a nominal volumetric concentration of 16% (v/v) in diethyl phthalate and further diluted 8 times with air for a final concentration of 2% for epifluorescence imaging. For multiphoton imaging odors were diluted in mineral oil at 16% (v/v) and diluted 16 times with air for a final concentration of 1%. For most experiments, odors were presented for 5s with an interstimulus interval of at least 40s.

The odor panel for epifluorescence imaging consisted of 1) Methyl propionate 2) Methyl butyrate 3) Ethyl Valerate 4) Pentyl acetate 5) Propyl acetate 6) Valeraldehyde 7) Methyl tiglate.

The odor panel for multiphoton imaging consisted of 1) Ethyl tiglate 2) Ethyl valerate 3) Valeric acid 4) Allyl butyrate 5) Isoamylamine 6) 2-Methoxypyrazine 7) Isugenol 8) Methyl tiglate 9) Valeraldehyde 10) Isoamyl acetate 11) Heptanal 12) mineral oil 13) Ethyl propionate 14) Propyl acetate 15) 2-Heptanone 16) Acetophenone.

For imaging glomerular responses to odor concentrations an additional 16 channel olfactometer outfitted with two odors, Ethyl valerate and Allyl butyrate, was used. The initial concentration series for each odor was 80%, 16%, 8%, 1.6%, 0.8%, 0.16%, 0.08% (v/v) in mineral oil and further diluted 16 times with air. Odors were presented for 2s to prevent adaptation at the strongest concentrations. For all experiments, odors were delivered 3-5 times each.

#### Analysis

Calcium signals were extracted from raw images using custom-written scripts in MATLAB (MathWorks Inc.) and reported as ΔF/F signals, where F represents the average baseline fluorescence. Regions of interest were selected from average fluorescence projections for multiphoton imaging and ΔF/F projections for epifluorescence imaging. Response amplitude was measured from between three and five repeats of each odor as the mean response in the 5 seconds following odor onset. Bleaching was corrected by fitting a single exponential to blank odor trials in multiphoton imaging and fitting a single exponential to the baseline period for epifluorescence experiments. For images of ΔF/F signals, the mean of an equal number of median filtered frames in the baseline and odor period was used. Traces of ΔF/F signals were smoothed for display. For figures where a threshold was applied to the data, thresholds were calculated based on the distribution of blank odor trials. An area under the receiver operating curve analysis was performed and the lowest threshold yielding ten responses for every one blank odor response was chosen. Sparseness measures were calculated as previously reported^48,49^. Statistical comparisons for imaging experiments were made as described in the text for each figure and values are given as mean +/-standard error of the mean.

### Respiration measurements

#### Surgery

Animals were anesthetized with ketamine/xylazine as described above and a head plate was implanted in the skull as described previously in this article. For some mice, a small craniotomy was also made through the right nasal bone (1 mm anterior from the frontal/nasal fissure, 1mm lateral from the midline), and a hollow cannula (#C313G; Plastics One Inc.) was lowered into the hole and glued to the skull. Finally, the whole exposed skull was covered with dental cement (Jet Repair, Lang Dental). The mice were given a week after the surgery to recover before any experiment was performed.

#### Respiration monitoring

Two strategies were used to monitor the breathing: measuring the intranasal pressure through an implanted cannula^50,51^, and measuring the temperature in front of the nose^52^.

For the intranasal pressure strategy, mice previously implanted with a cannula were head-fixed. Then, the cannula was connected to a pressure sensor (24PCEFJ6G; Honeywell International) through a piece of polyethylene tubing. The voltage signal generated by the sensor was amplified 1000x, low-pass filtered at 60Hz, and digitized at 1000Hz using custom software written in Labview.

For the temperature measurement, mice were head-fixed, and a thermocouple (5TC-TT-JI-40-1M, Omega Engineering) was placed ~ 2mm in front of their nose. The voltage changes generated by the temperature variations were amplified 10000x, low-pass filtered at 60Hz, and digitized at 1000Hz using custom software written in Labview.

#### Analysis

Analysis of breathing signals and statistical tests were performed using custom software written in MATLAB. Two types of statistical tests were used: the Kruskal-Wallis test, and a bootstrap test to compare the means of two distributions (MATLAB function “bootstrp()” repeated 1,000,000 times for each bootstrapped statistics). The MATLAB toolbox CircStat was used to analyze circular data^53^. The critical p-value was set at 5% for all the tests, and Bonferroni correction was applied for multiple comparisons. On the figures, all the values are given as mean +/-standard error of the mean, unless otherwise stated.

### Open Field Behavior

The arena consisted of 56cm diameter circular inner chamber with four air inlets equally spaced around its circumference. The circular inner chamber was housed in light- and sound-proof outer chamber and illuminated using infrared LEDs. Throughout each experiment, airflow was maintained at a constant velocity for each inlet. After 10 minutes baseline exploration, air to one of the inlets was redirected through an odorized chamber while ensuring no change in its velocity. A vacuum was located at the center of the arena and its flow matched to the sum of all air inlets to prevent the accumulation of odor in the arena. Peanut oil was diluted in mineral oil as in imaging experiments. Each mouse was only tested once and the order in which they were tested was randomized. After each mouse the arena was thoroughly cleaned with ethanol to eliminate the presence of social cues. Images for tracking were acquired at 8Hz using a USB camera (Grasshopper3, Point Grey Imaging) and custom-written Labview software. Images were processed using custom MATLAB routines to measure location and velocity. Mice with an initial position >10cm from the odor source were excluded from our analysis. All statistical comparisons for behavior experiments were made with Wilcoxon rank-sum test and values are given as mean +/-standard error of the mean.

## Author Contributions

JDZ and VNM designed the research. JDZ and JG collected and analyzed the data. R-CL and C-CL characterized and genotyped the ANO2-null mouse. JDZ and VNM wrote the manuscript with input from all authors.

## Acknowledgements

We thank Vikrant Kapoor for technical assistance. We also thank members of the Murthy Lab and Prof. King-Wai Yau for helpful discussions. JDZ, JG, and VNM were partly supported by a grant from the NIH (R01 DC14454). JDZ was supported by NIH Fellowship F32 DC015938. R-CL and C-CL were supported by a grant from the NIH (R01 DC14941 to King-Wai Yau).

---

**Supplemental Figure S1:**
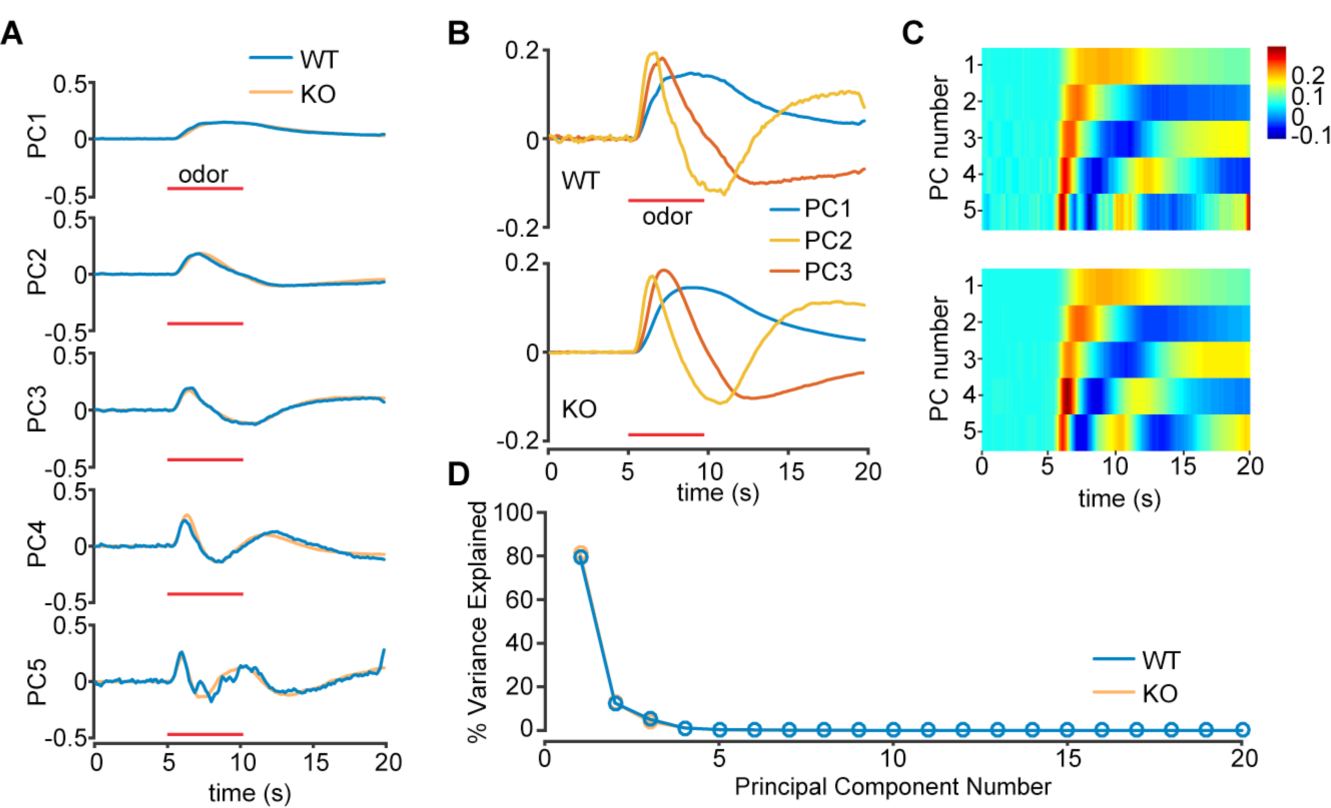
Principal component analysis of ORN responses measured using epifluoresence microscopy. **A**. Overlaid time course of the first five principal components of all responses above threshold for WT and KO animals. Red bar indicates odor delivery period. **B**. The first three principal components combined and plotted on the same axis. **C**. The first five principal components combined and plotted on the same axis. **D**. Plot of the variance explained by each of the first 20 principal components.

**Supplemental Figure S2:**
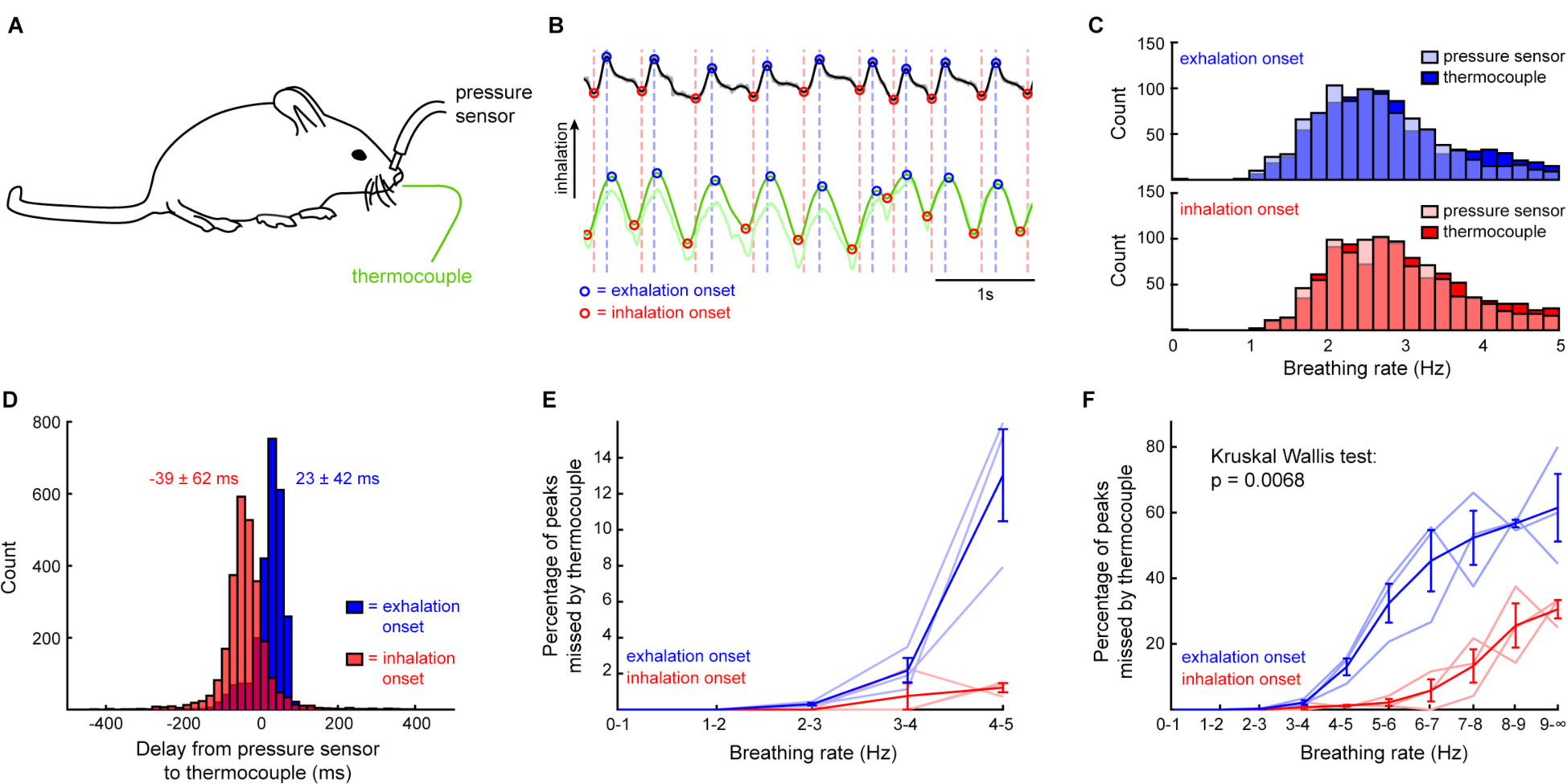
Validation of the external thermal sensor versus pressure sensor for respiration monitoring. **A**. Experimental setup. The breathing of head-fixed, awake mice (n = 3) was monitored through two methods: a cannula implanted in the nasal cavity, connected to a pressure sensor, and an external thermal sensor (thermocouple) in front of the nostrils. **B**. Examples of respiration traces. Inhalation is upward. Light traces show raw data and dark traces are the data after digital filtering. Black: pressure sensor. Green: thermocouple. The blue dots show the peaks of inhalation, while the red ones show the peaks of exhalation. **C**. Histograms of the instantaneous respiration rate of an exemplar mouse using a pressure sensor (darker color) or the thermocouple (lighter color). Inhalation is the histogram on top in blue and exhalation is the histogram in red on bottom. **D**. Histogram of the delay between peak inhalation and exhalation measured from the pressure sensor and thermocouple across all respirations. Values next to each distribution: mean +/-standard deviation. **E**. Fraction of the inhalation or exhalation peaks missed by the thermocouple, as a function of the breathing rate. Change this Each lighter curve is a different mouse. Each darker curve gives the mean +/-standard error of the mean. At respiration frequencies lower than 5Hz, which is typical of anesthetized mice, the thermocouple is a reliable method for the monitoring of breathing rates **F**. Graph from part E expanded to higher respiration rates. At respiration frequencies higher than 5Hz, which is only seen in awake mice, the thermocouple is not a reliable method for the monitoring of breathing rates (Kruskal Wallis test to compare the two distributions, p = 0.007).

**Supplemental Figure S3:**
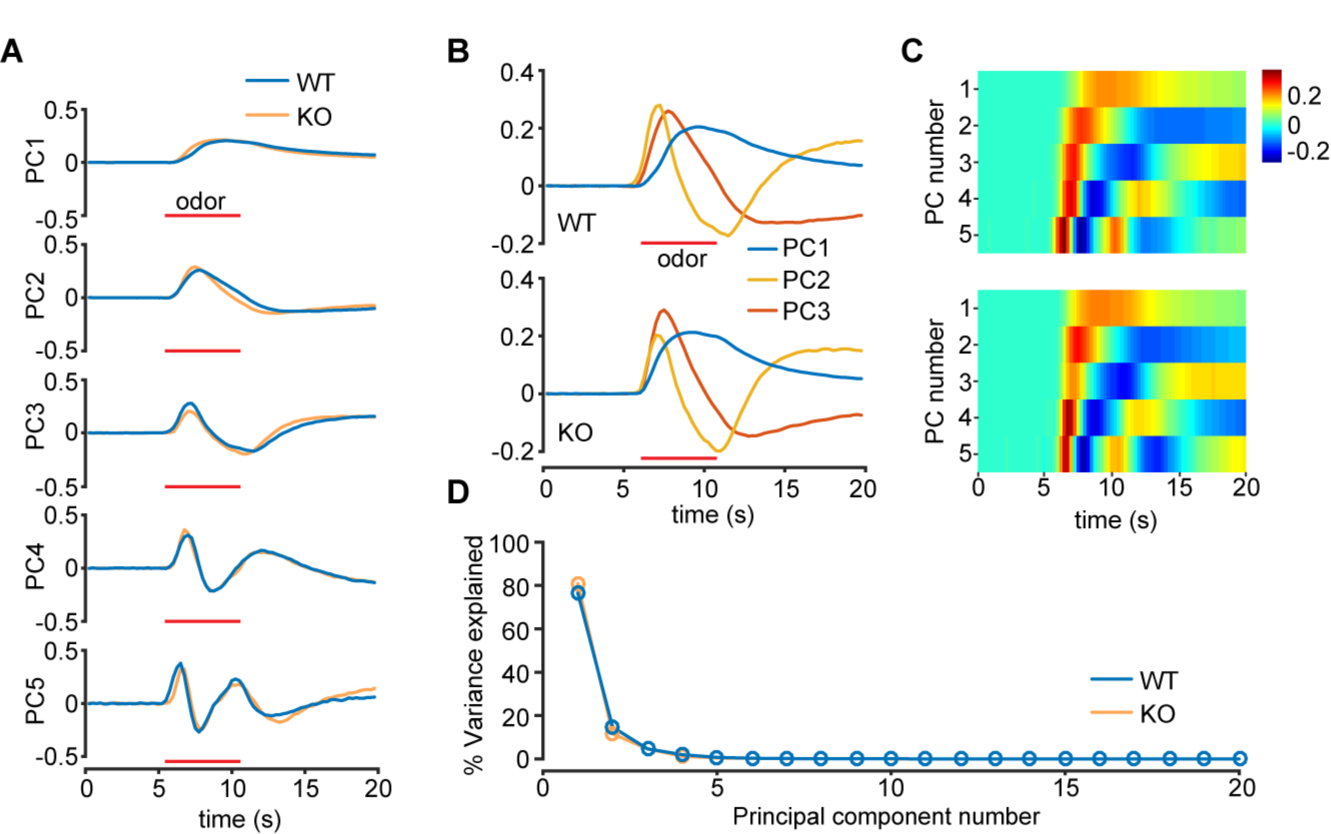
Principal component analysis of ORN responses using multiphoton microscopy. **A**. Overlaid time course of the first five principal components of all responses above threshold for WT and KO animals. Red bar indicates odor delivery period. **B**. The first three principal components combined and plotted on the same axis. **C**. The first five principal components combined and plotted on the same axis. **D**. Plot of the variance explained by each of the first 20 principal components.

**Supplemental Figure S4:**
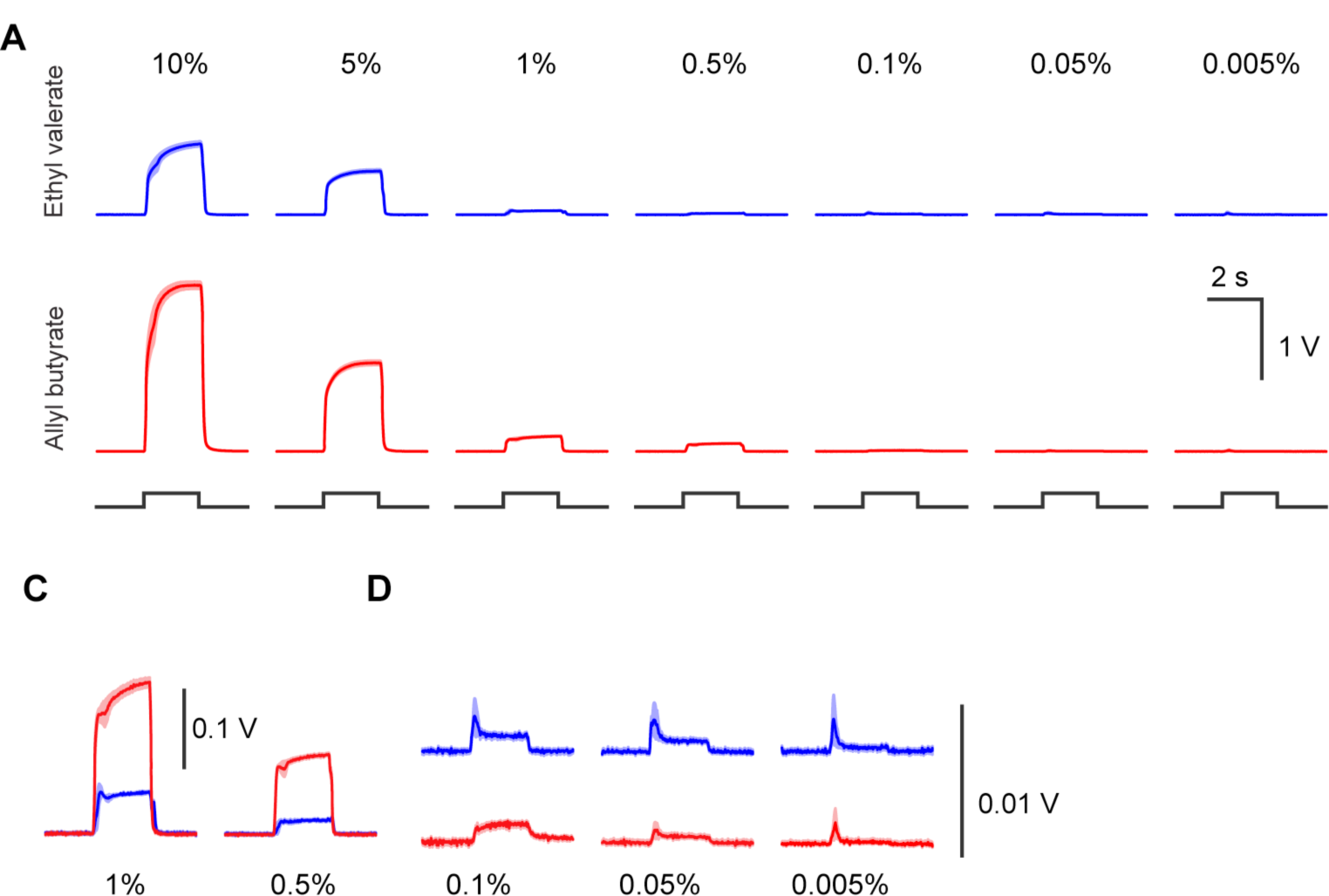
Photoionization detector measurements of odor concentrations. **A**. Mean photoionization detector (PID) signals across five repeats for Ethyl valerate and Allyl butyrate. Shaded area each trace is the s.e.m. Black line below indicates the digital signal used to open each solenoid. **B**. Expanded and overlaid PID signals from odors measured at 1% and 0.5%. **C**. Expanded PID signals from odors at 0.1%, 0.05%, and 0.005%. Signals are expanded 100 times from part A.

